# Systematic determination of the mitochondrial proportion in human and mice tissues for single-cell RNA sequencing data quality control

**DOI:** 10.1101/2020.02.20.958793

**Authors:** Daniel Osorio, James J. Cai

**Affiliations:** Department of Veterinary Integrative Biosciences, Texas A&M University, College Station, TX 77843-4458, USA; Department of Electrical & Computer Engineering, Texas A&M University, College Station, TX 77843-4458, USA; Interdisciplinary Program of Genetics, Texas A&M University, College Station, TX 77843-4458, USA

## Abstract

**Motivation:** Quality control (QC) is a critical step in single-cell RNA-seq (scRNA-seq) data analysis. Low-quality cells are removed from the analysis during the QC process to avoid misinterpretation of the data. One of the important QC metrics is the mitochondrial proportion (mtDNA%), which is used as a threshold to filter out low-quality cells. Early publications in the field established a threshold of 5% and since then, it has been used as a default in several software packages for scRNA-seq data analysis and adopted as a standard in many scRNA-seq studies. However, the validity of using a uniform threshold across different species, single-cell technologies, tissues, and cell types has not been adequately assessed.

**Results:** We systematically analyzed 5,530,106 cells reported in 1,349 annotated datasets available in the PanglaoDB database and found that the average mtDNA% in scRNA-seq data across human tissues is significantly higher than in mouse tissues. This difference is not confounded by the platform used to generate the data. Based on this finding, we propose new reference values of the mtDNA% for 121 tissues of mice and 44 tissues of humans. In general, for mouse tissues, the 5% threshold performs well to distinguish between healthy and low-quality cells. However, for human tissues, the 5% threshold should be reconsidered as it fails to accurately discriminate between healthy and low-quality cells in 29.5% (13 of 44) tissues analyzed. We conclude that omitting the mtDNA% QC filter or adopting a suboptimal mtDNA% threshold may lead to erroneous biological interpretations of scRNA-seq data.

*Availability:* The code used to download datasets, perform the analyzes, and produce the figures is available at https://github.com/dosorio/mtProportion

*Contact:*

**Supplementary information:** Supplementary data are available at *Bioinformatics* online.

## 1 Introduction

Single-cell RNA-seq (scRNA-seq) experiments have improved the resolution of our knowledge about cellular composition and cellular behavior in complex tissues (Sandberg 2014). A critical step during scRNA-seq data processing is to perform quality control (QC) over the cells sequenced transcriptomes (Hwang et al. 2018). The QC process usually involves applying user-defined thresholds for different metrics computed for each individual cell to filter out doublets and “low-quality” cells (Luecken and Theis 2019). Commonly used QC metrics include the total transcript counts (also known as the library size), the number of expressed genes, and the mitochondrial proportion (mtDNA%, i.e., the ratio of reads mapped to mitochondrial DNA-encoded genes to the total number of reads mapped). Defining the proper thresholds of QC metrics is a complex task that requires a vast knowledge of the cellular diversity in the tissue under study. Thresholds may be uniquely set for each sample, as they are dependent on the cells or tissue being processed (Ji and Sadreyev 2019).

Our present study focuses on the systematic determination of a threshold for mtDNA%—the fraction of mitochondrial counts per cell—in scRNA-seq QC. Mitochondrial content is known to interact with the nuclear genome, drive alternative splicing, and regulate nuclear gene expression, and is also associated with cancer, degenerative diseases, and aging (Guantes et al. 2015; Muir et al. 2016). High numbers of mitochondrial transcripts are indicators of cell stress, and therefore mtDNA% is a measurement associated with apoptotic, stressed, low-quality cells (Zhao et al. 2002; Ilicic et al. 2016; Lun et al. 2016). However, mtDNA% threshold depends highly on the tissue type and the questions being investigated (AlJanahi et al. 2018). The mtDNA% threshold is of economic and biological importance. A wrongly defined, very stringent mtDNA% threshold may cause bias in the recovered cellular composition of the tissue under study. This bias may force the researchers to increase the sample size to capture enough cells (which may not have the normal biological behavior of the cell type) under the threshold, and thus increase the cost of the experiment. Inversely, a relaxed threshold of mtDNA% may allow apoptotic, low-quality cells to remain in the analysis, resulting in the identification of wrong biological patterns.

To reduce the bias caused by the use of arbitrary mtDNA% thresholds, Ma and collaborators (Ma et al. 2019) proposed an unsupervised method to optimize the threshold for each given input data. This computationally expensive data-driven procedure, which defines the threshold as a function of the distribution of the data, due to the lack of reference values, is not able to identify bias induced during the library preparation. Without such standard references, the values of mtDNA% thresholds fluctuate with different input data sets. For example, a largely failed experiment may generate a data set, in which most cells have an inflated mtDNA%. Accordingly, the optimized threshold based on these inflated values may be unreasonably high. Therefore, having a uniform and standardized threshold for scRNA-seq data analysis is essential. It improves the reproducibility of experiments and simplifies the automatization of bioinformatic pipelines (McCarthy et al. 2017).

Through analysis of bulk RNA-seq data produced by the Illumina Body Tissue Atlas, Mercer and collaborators reported the mtDNA% for 16 human tissues (Mercer et al. 2011). They found that the mtDNA% ranges from 5% or less in tissues with low energy requirements up to approximately 30% in the heart due to the high energy demand of cardiomyocytes. Based on that study, early publications of scRNA-seq datasets used the 5% threshold reported for tissues with low energy demands (e.g., adrenal, ovary, thyroid, prostate, testes, lung, lymph, and white blood cells) as default for data quality control (Lukassen et al. 2018). Furthermore, the 5% threshold has been adapted as the default parameter by Seurat—one of the most popular software packages for scRNA-seq data analysis (Satija et al. 2015). These have made 5% a practical standard for scRNA-seq data analyses. Nevertheless, due to the lack of reference values for mtDNA% in different species, technologies, tissues and cell types, the optimal value for a standardized threshold is still an open question in the field.

PanglaoDB is a scRNA-seq database providing uniformly processed, annotated count matrices for thousands of cells from hundreds of scRNA-seq experiments. The data source of PanglaoDB is the sequence read archive (SRA) database of the National Center for Biotechnology Information (NCBI). With the data sets from PanglaoDB, it is possible to systematically evaluate the optimal threshold of mtDNA% for different experimental settings that may vary across platforms, technologies, species, tissues or cell types (Franzen et al. 2019; Svensson et al. 2019). Here, we present a systematic analysis of the mtDNA% in more than 5 million cells reported in over one thousand data sets in PanglaoDB (Franzen et al. 2019). We compared the mtDNA% reported for different technologies, species, tissues, and cell types. By analyzing the data provided by hundreds of experiments together, we reach the consensus reference values for more than 40 human tissues and more than 120 mouse tissues. Furthermore, we evaluated the validity of using the 5% threshold in different humans and mice tissues and showed that omitting the mtDNA% as a QC filter led to erroneous biological interpretations of the data.

## 2 Methods

Datasets in the PanglaoDB database, available at the time of analysis (in January 2020), were downloaded and processed using R 3.6.2 (R Core Team 2013) through an ‘*in-house*’ script using the XML (Lang and CRAN Team 2012) and xml2 (Wickham et al. 2018) packages. The library size (total number of counts), the total number of detected genes, and the total number of counts that match with the mitochondrial genes (mitochondrial counts) were estimated for all cells in each of downloaded datasets. The SRA/SRS identifiers, species, protocol, tissue, cell type and barcode of each experiment were obtained and associated with each cell.

Only the cells with more than 1,000 counts and with the total number of counts greater than two times the average library size in the same sample were retained for analysis. In addition, a polynomic regression of degree 2 (to account for saturation) was applied to establish the 95% confidence intervals of the predicted total number of genes as a function of the library size per cell. Cells with an observed total number of genes below or above expectation limits were removed from the analysis. The same procedure was applied a second time to establish the 95% confidence intervals of the predicted mitochondrial counts as a function of library size. An ordinary least squares (OLS) regression model was used to fit the data and cells with exceptionally high or low mitochondrial counts were removed from the analysis.

Subsequently, the mtDNA% value was computed for each cell as the ratio between the mitochondrial counts and the library size of the cell. The mtDNA% values were then compared between cells from different settings: species, technologies, tissues, and cell types. To compare the mtDNA% between humans and mice cells, we used the Welch two-sample *t*-test and used the Wilcoxon sum-rank test to cross-validate the results. To evaluate the reliability of the 5% threshold, a comparison to evaluate whether the mean was less than the 0.05 threshold value was performed using the t-test for each tissue and cell type independently using the data generated by the 10x Genomics Chromium system, after filtering out groups with less than 1,000 cells.

Example datasets (SRS3703557, SRS3545826, and SRS2397417) were downloaded from the PanglaoDB along with cell clustering results and cell type information. Count matrices were processed using the ‘Seurat’ R package to generate low dimensional representations. Differential expression analysis was performed using ‘MAST’ (Finak et al. 2015) to compare the transcriptome profile of clusters exhibiting a median mtDNA% higher than 5% against that of other clusters with the same cell type but with a median mtDNA% lower than 5%. By using the sorted list of fold-changes reported by MAST, we performed Gene Set Enrichment Analysis (GSEA) to test the enrichment of the ‘*Apoptosis*’ pathway from the KEGG database (Kanehisa and Goto 2000) in the clusters with increased mtDNA%. To run the GSEA analysis efficiently, we used the multilevel function included in the ‘fgsea’ R package (Korotkevich et al. 2019).

## 3 Results

We downloaded a total of 5,530,106 cells reported in 1,349 datasets from the PanglaoDB database. From those, we removed 278,607 cells with a total number of counts smaller than 1,000 or above two times the average library size in the sample where it was sequenced. Also, 80,225 cells with no mitochondrial counts were removed. The remaining 5,171,274 cells were used to establish the 95% confidence intervals of the predicted total number of genes as a function of the library size per cell (Fig. S1). We found that the relationship between the number of genes and the library size is monotonically positive (*ρ* = 0.89; *P* < 2.2 × 10^−16^), which is consistent with that previously reported (Svensson et al. 2019). We also found that the expected total number of genes reaches saturation at a point close to the 1 × 10^5^ library size counts. In this step, we removed 157,960 cells because they have a total number of quantified genes above (*n* = 5,509) or below (*n* = 152,451) the 95% confidence interval limit defined from the prediction.

Next, we accounted for outliers in the mitochondrial counts in relative to the library size. This procedure has been shown to be critical to differentiate apoptotic cells of pre-apoptotic and healthy cells in a supervised experiment (Ordonez-Rueda et al. 2020). To do so, we used the OLS regression and computed the confidence interval of prediction between the mitochondrial counts and the library size with data from all 5,013,314 cells. We found that the relationship is noisy but positive and linear (Fig. S2; *r* = 0.65, *P* < 2.2 × 10^−16^). Following this procedure, we identified 333,712 cells with mitochondrial counts above (*n* = 178,671) or below (*n* = 1 55,041) the computed confidence interval limits, which were also removed. After this step, 4,679,602 cells were retained for the study.

With the cleaned dataset, we estimated that the mtDNA% per cell is distributed between the minimum of 0.17% and the maximum of 14.64%, considerably lower than the upper limit previously reported (up to 30% in heart) using the bulk RNA-seq generated by the Illumina Body Tissue Atlas (Mercer, Neph et al. 2011). Next, we performed a comparison to evaluate whether there is a difference in the average mtDNA% cross different species. The PanglaoDB database contains human and mouse datasets; therefore, our comparison was between human and mouse. We performed the Welch two-sample t-test and use the Wilcoxon-sum rank test to validate the results. Both tests converged to the same conclusion, that is, the average mtDNA% in human cells is significative higher than that in mice cells (*P* < 2.2 × 10^−16^, in both cases) as is displayed in Fig. 1A.

**Fig 1.**
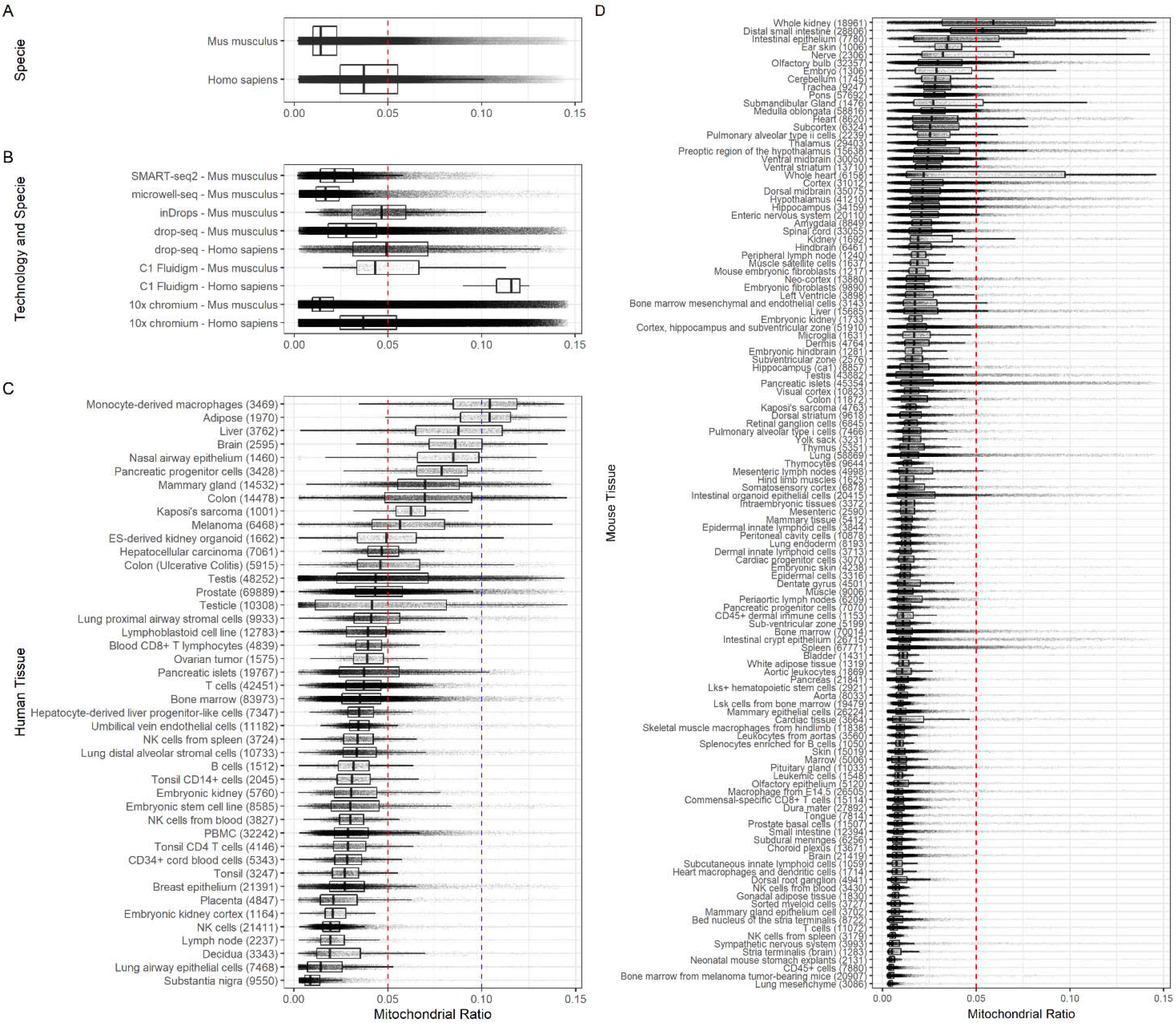
Boxplots showing the differences in mtDNA% across species, technologies and tissues. Each dot represents a cell; the red line is the early established 5% threshold, and the blue line is the 10% threshold for human cells proposed here. In parenthesis (panel C and D), the number of cells in the stated tissue. **(A)** The difference in mtDNA% between human and mice cells. **(B)** The differences in mtDNA% between human and mice cells by the technology used to generate the data. **(C)** Boxplots of mtDNA% across 44 human tissues. **(D)** Boxplots of mtDNA% across 121 mouse tissues.

Then, we compared the mitochondrial content between human and mouse data, stratified by the type of scRNA-seq technologies, by which the data is obtained. These technologies include drop-seq, C1 Fluidigm and 10x Genomics. Our results confirm our previous finding. In all cases wherever data allowed, no matter which technology is used, the same pattern was recovered. That is, Human cells have significantly larger mtDNA% than mice cells (Fig. 1B). Most importantly, for all cases where mitochondrial content in humans was evaluated, the 75th percentile was located above the threshold, suggesting that the early defined 5% is not appropriate for human cells. Note that 91.3% (*n* = 4,271,613) of cells analyzed here were processed using the 10x Genomics chromium system. Next, we decided to perform the comparison of the mitochondrial content between tissues and cell types using only the data generated using the 10x Genomics technology.

For humans, we identified 44 tissues, for which more than 1,000 cells are available in the database. From those 44 tissues, 13 (29.5%) showed an average mtDNA% significantly higher than 5%. The 13 human tissues are nasal airway epithelium, monocyte-derived macrophages, testicle, colon (ulcerative colitis), liver, colon, melanoma, mammary gland, ES-derived kidney organoid, pancreatic progenitor cells, adipose, Kaposi’s sarcoma, and brain. However, as displayed in Fig. 1C, 18 of the 44 human tissues (40%) have a portion of the interquartile range over the 5% threshold. Only two of them, monocyte-derived macrophages and adipose, have an average mtDNA% higher than 10%. This result supports our observation that the early defined 5% is not appropriate for human tissues. We conclude that the new standardized threshold for human tissues should be 10% instead. At the cell-type level, we found similar patterns. From 37 different cell types with more than 1,000 cells derived from human samples, 13 of them (35.1%) have an average mtDNA% greater than 5%, but none of them have an average mtDNA% greater than 10% (Fig. S3). The 13 cell types are hepatocytes, epithelial cells, neutrophils, cholangiocytes, smooth muscle cells, keratinocytes, Langerhans cells, spermatocytes, ductal cells, beta cells, luminal epithelial cells, macrophages, and embryonic stem cells. Furthermore, only 4 of them (epithelial cells, Langerhans cells, spermatocytes, and macro-phages) have a portion of the interquartile range above the 10% threshold. For mice, when the mtDNA% was compared across the cell types with at least 1000 cells reported in the database, 7 of 74 cell types showed an average mtDNA% greater than 5%. The 7 cell types are proximal tubule cells, distal tubule cells, hepatocyte, cardiomyocytes, Leydig cells, intercalated cells, choroid plexus cell (Fig S4). In contrast to the identified 44 human tissues, there are many more mouse tissues (121) with more than 1,000 cells reported in the database. Among them, only 3 (2.5%) showed an average mtDNA% significantly higher than 5% (whole kidney, whole heart, and distal small intestine). Furthermore, only 6 mouse tissues (whole kidney, intestinal epithelium, whole heart, nerve, distal small intestine, and submandibular gland) have a portion of the interquartile range over the 5% threshold (Fig. 1D). These findings indicate that the 5% threshold early proposed in the field is an appropriate standardized threshold for mouse tissues.

To evaluate the effect of mtDNA% in the analysis of single-cell RNA-seq data, we downloaded three datasets from the PanglaoDB database using accessions: SRS3703557, SRS3545826, and SRS2397417. The first dataset contains 9,238 cells from the mouse heart, the second 7448 cells from mouse lung, and the third 9,057 cells from the human umbilical vein.

First, we evaluated the effect of genes encoded in the mitochondrial genome (for short, mtGenes) on the cell clustering results. We compared the low dimensional representations generated by t-SNE using the PCA result as a prior with and without the mtGenes. We found that, in all three examples, the mitochondrial content does not affect significantly the structure of the data in low dimensional representation, allowing to recover clearly in both cases (with and without mtGenes) the clusters reported by the PanglaoDB database (Fig. 2 and Fig. S5). These results confirm findings previously reported in an evaluation study of different computational pipelines for scRNA-seq data pre-processing (Germain et al. 2020). We also found that even without taking into account the mtGenes for the generation of the low dimensional representation, low-quality cells with high mtDNA% tend to cluster together (Fig. 2B; Clusters 19, 0, 9, 7, and 11 in SRS3703557; cluster 5 in SRS3545826, and cluster 11 in SRS2397417).

**Fig 2.**
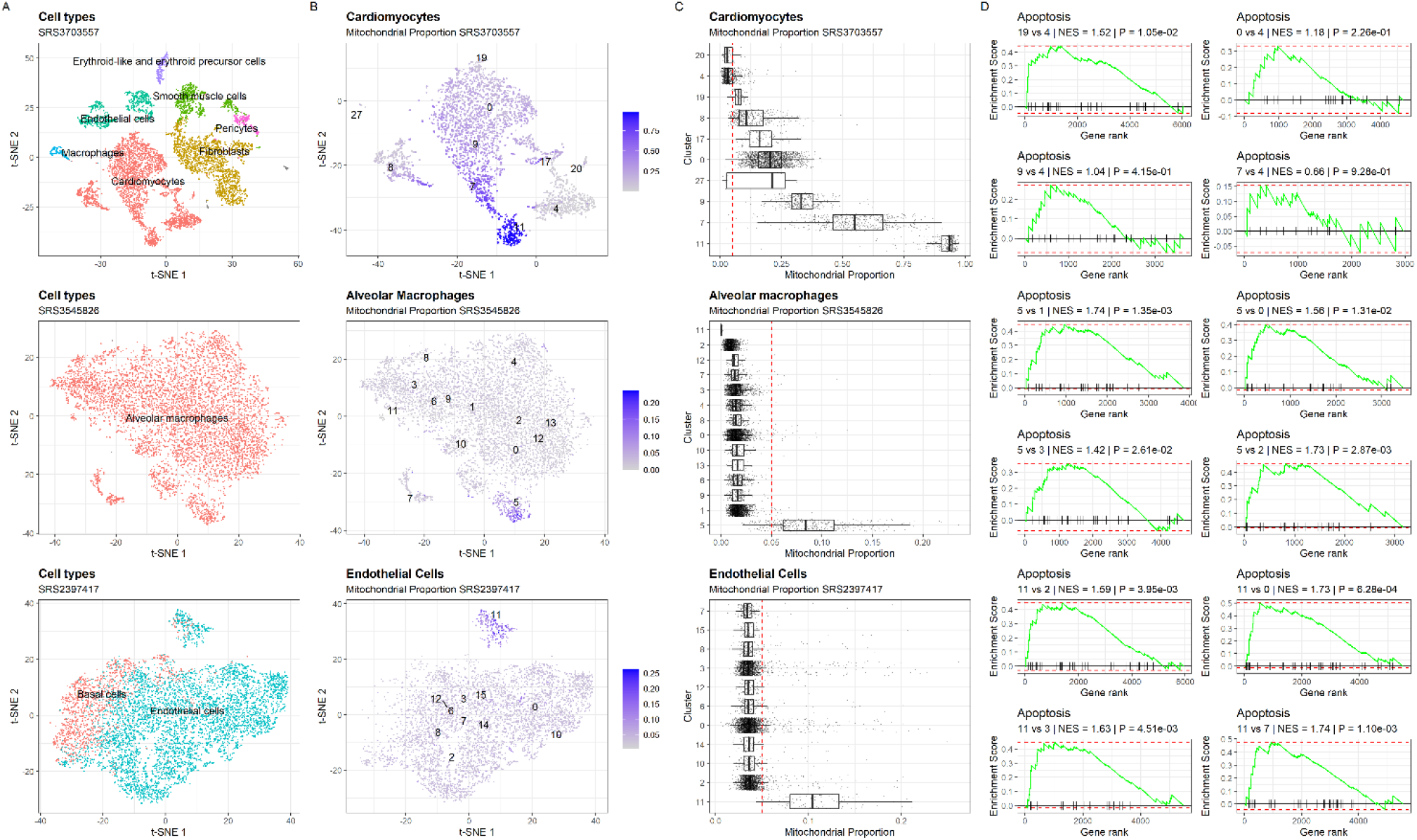
Case examples showing the effect of omitting the mtDNA% QC filter in the analysis of scRNA-seq data. **(A)** t-SNE representation of all the cell populations included in the dataset generated by excluding the mitochondrial genes from the list of highly variable genes before principal component analysis (PCA). Each dot represents a cell and they are colored by cell-type. **(B)** t-SNE representation of cell-type used as an example colored in the function of the mtDNA% in each cell. Clusters reported by the PanglaoDB are labeled. **(C)** Boxplot showing the distribution of the mtDNA% across clusters. The red line is the early established 5% threshold. **(D)** GSEA analysis of the *Apoptosis* pathway between clusters with a high proportion of low-quality cells and others containing high-quality cells.

Next, we evaluated the significance of the threshold to identify low quality cells. For each example, after identifying the clusters containing cells with high mtDNA% of a cell type, we used MAST to compare their expression profiles against other clusters with most of the cells below the threshold. The fold-change values reported by MAST were used as the input of GSEA analysis to test for the significance of the *Apoptosis* pathway in the KEGG database.

For the first example, we focused on cardiomyocytes, a cell type associated with high mtDNA% with nine clusters reported by the PanglaoDB database (Fig. 2C) for this dataset. We compared cells with a higher mtDNA% level in clusters 19, 0, 9, 7, and 11, which form a larger cluster, against cells with a lower mtDNA% level in cluster 4. We found that the number of genes with detectable expression decreases with the increase of mtDNA% in cells (as shown by the x-axes of the first two rows of subplots in Fig. 2D). This anticorrelation is expected as the increased mtDNA% is likely to be associated with cell breakout events. When a breakout occurs (due to the differences in copy numbers given by a single nucleus and several mitochondria by cell), generate an increase in the abundance of mitochondrial content, and reducing the reads that will be mapped to nuclear genes, resulting in fewer genes’ expression detected. Indeed, we found even a small increase of the mitochondrial content (comparing cluster 19 and cluster 0 vs. cluster 4) led to a huge decrease in the number of expressed genes (>6,000 genes vs. ~5,000 genes, see the first row of subplots in Fig. 2D). The number of genes included in the GSEA analysis, in turn, influences the value of Normalized Enrichment Score (NES), which is used to assess the significance of the apoptosis pathway. Despite this influence, a positive NES value was recovered in all the cases for the tested cardiomyocytes clusters, as well as in the other two examples of mouse alveolar macrophages (cluster 5 against others) and human endothelial cells (cluster 11 against others), suggesting a consistently higher expression (positive log_2_ fold-change) of apoptotic pathway genes among cells with mtDNA % above the threshold (Fig. 2D).

In summary, we reported a new set of mtDNA% reference values across human and mice tissues and cell-types for scRNA-seq QC (Table S1-S4). Based on our analytical results, we suggest a standardized mtDNA% threshold of 10% for scRNA-seq QC of human samples. For mouse samples, we found that the early defined threshold of 5% accurately discriminates between healthy and low-quality cells, bringing to evidence that under a well-performed scRNA-seq QC, clusters containing cells with high mtDNA% exhibiting signatures of apoptosis, like those shown in the example datasets, should be excluded from being used to make biological interpretations. Thus, we suggest that all published mouse studies, in which scRNA-seq QC was based on the mtDNA% value greater than 5%, should be reevaluated because the use of any mtDNA% higher than 5% is likely to be an overshoot over the threshold, resulting in apoptotic cells being utilized in the subsequent analyses.

## Acknowledgements

We thank Dr. Chapkin and lab members from the Texas A&M nutrition department for their thoughtful questions that motivated this paper.

D.O. was supported by the 2020 Award of Texas A&M Institute of Data Science (TAMIDS) Data Resource Development Program.

## References

AlJanahi AA, Danielsen M, Dunbar CE. 2018. An Introduction to the Analysis of Single-Cell RNA-Sequencing Data. Mol Ther Methods Clin Dev 10: 189–196.

Finak G, McDavid A, Yajima M, Deng J, Gersuk V, Shalek AK, Slichter CK, Miller HW, McElrath MJ, Prlic M. 2015. MAST: a flexible statistical framework for assessing transcriptional changes and characterizing heterogeneity in single-cell RNA sequencing data. Genome biology 16: 1–13.

Franzen O, Gan LM, Bjorkegren JLM. 2019. PanglaoDB: a web server for exploration of mouse and human single-cell RNA sequencing data. Database (Oxford) 2019.

Germain P-L, Sonrel A, Robinson MD. 2020. pipeComp, a general framework for the evaluation of computational pipelines, reveals performant single-cell RNA-seq preprocessing tools. BioRxiv.

Guantes R, Rastrojo A, Neves R, Lima A, Aguado B, Iborra FJ. 2015. Global variability in gene expression and alternative splicing is modulated by mitochondrial content. Genome Res 25: 633–644.

Hwang B, Lee JH, Bang D. 2018. Single-cell RNA sequencing technologies and bioinformatics pipelines. Exp Mol Med 50: 96.

Ilicic T, Kim JK, Kolodziejczyk AA, Bagger FO, McCarthy DJ, Marioni JC, Teichmann SA. 2016. Classification of low quality cells from single-cell RNA-seq data. Genome Biol 17: 29.

Ji F, Sadreyev RI. 2019. Single-Cell RNA-seq: Introduction to Bioinformatics Analysis. Curr Protoc Mol Biol 127: e92.

Kanehisa M, Goto S. 2000. KEGG: kyoto encyclopedia of genes and genomes. Nucleic acids research 28: 27–30.

Korotkevich G, Sukhov V, Sergushichev A. 2019. Fast gene set enrichment analysis. BioRxiv: 060012.

Lang DT, CRAN Team. 2012. XML: Tools for parsing and generating XML within R and S-Plus. 3.9–4.1.

Luecken MD, Theis FJ. 2019. Current best practices in single-cell RNA-seq analysis: a tutorial. Mol Syst Biol 15: e8746.

Lukassen S, Bosch E, Ekici AB, Winterpacht A. 2018. Single-cell RNA sequencing of adult mouse testes. Sci Data 5: 180192.

Lun AT, McCarthy DJ, Marioni JC. 2016. A step-by-step workflow for low-level analysis of single-cell RNA-seq data with Bioconductor. F1000Res 5: 2122.

Ma A, Zhu Z, Ye M, Wang F. 2019. EnsembleKQC: An Unsupervised Ensemble Learning Method for Quality Control of Single Cell RNA-seq Sequencing Data. In International Conference on Intelligent Computing, pp. 493–504. Springer.

McCarthy DJ, Campbell KR, Lun AT, Wills QF. 2017. Scater: pre-processing, quality control, normalization and visualization of single-cell RNA-seq data in R. Bioinformatics 33: 1179–1186.

Mercer TR, Neph S, Dinger ME, Crawford J, Smith MA, Shearwood AM, Haugen E, Bracken CP, Rackham O, Stamatoyannopoulos JA et al. 2011. The human mitochondrial transcriptome. Cell 146: 645–658.

Muir R, Diot A, Poulton J. 2016. Mitochondrial content is central to nuclear gene expression: Profound implications for human health. Bioessays 38: 150–156.

Ordonez-Rueda D, Baying B, Pavlinic D, Alessandri L, Yeboah Y, Landry JJM, Calogero R, Benes V, Paulsen M. 2020. Apoptotic Cell Exclusion and Bias-Free Single-Cell Selection Are Important Quality Control Requirements for Successful Single-Cell Sequencing Applications. Cytometry A 97: 156–167.

R Core Team. 2013. R: A language and environment for statistical computing.

Sandberg R. 2014. Entering the era of single-cell transcriptomics in biology and medicine. Nat Methods 11: 22–24.

Satija R, Farrell JA, Gennert D, Schier AF, Regev A. 2015. Spatial reconstruction of single-cell gene expression data. Nat Biotechnol 33: 495–502.

Svensson V, da Veiga Beltrame E, Pachter L. 2019. A curated database reveals trends in single-cell transcriptomics. bioRxiv.

Wickham H, Hester J, Ooms J. 2018. xml2: Parse XML. R Package Version 1.2.0.

Zhao Q, Wang J, Levichkin IV, Stasinopoulos S, Ryan MT, Hoogenraad NJ. 2002. A mitochondrial specific stress response in mammalian cells. EMBO J 21: 4411–4419.

