## Supplementary data for "Systematic determination of the mitochondrial proportion in human and mice tissues for single-cell RNA sequencing data quality control"

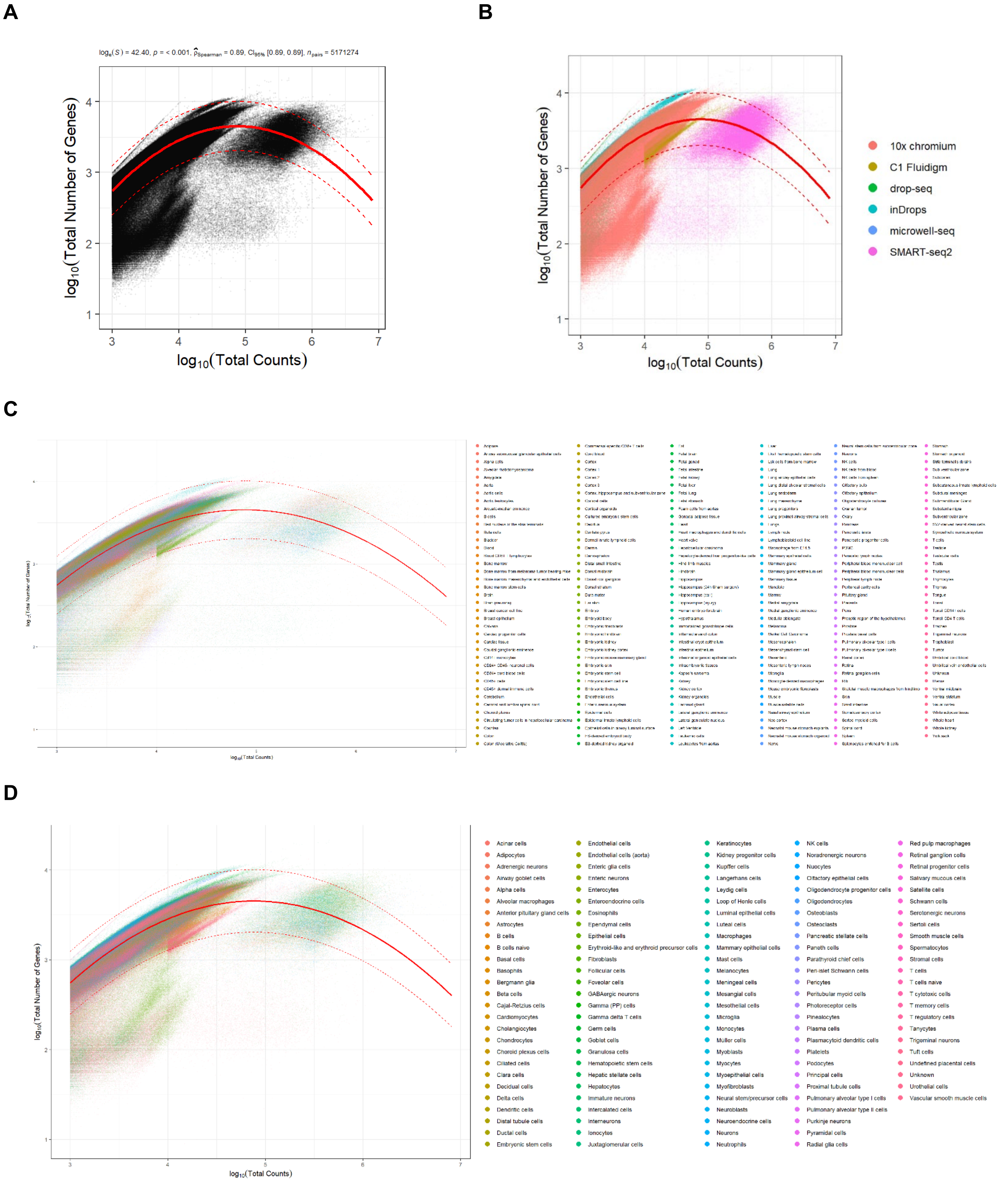


**Fig S1. Relationship between the total number of expressed genes and the library size.** (A) Each dot represents a cell, the continuous red line represents the expectation, and the dotted lines the lower and upper limit of the confidence interval for the prediction computed using a degree 2 regression. (B) Same as in A colored by technology. (C) Same as in A colored by tissue. (D) Same as in A colored by cell-type.

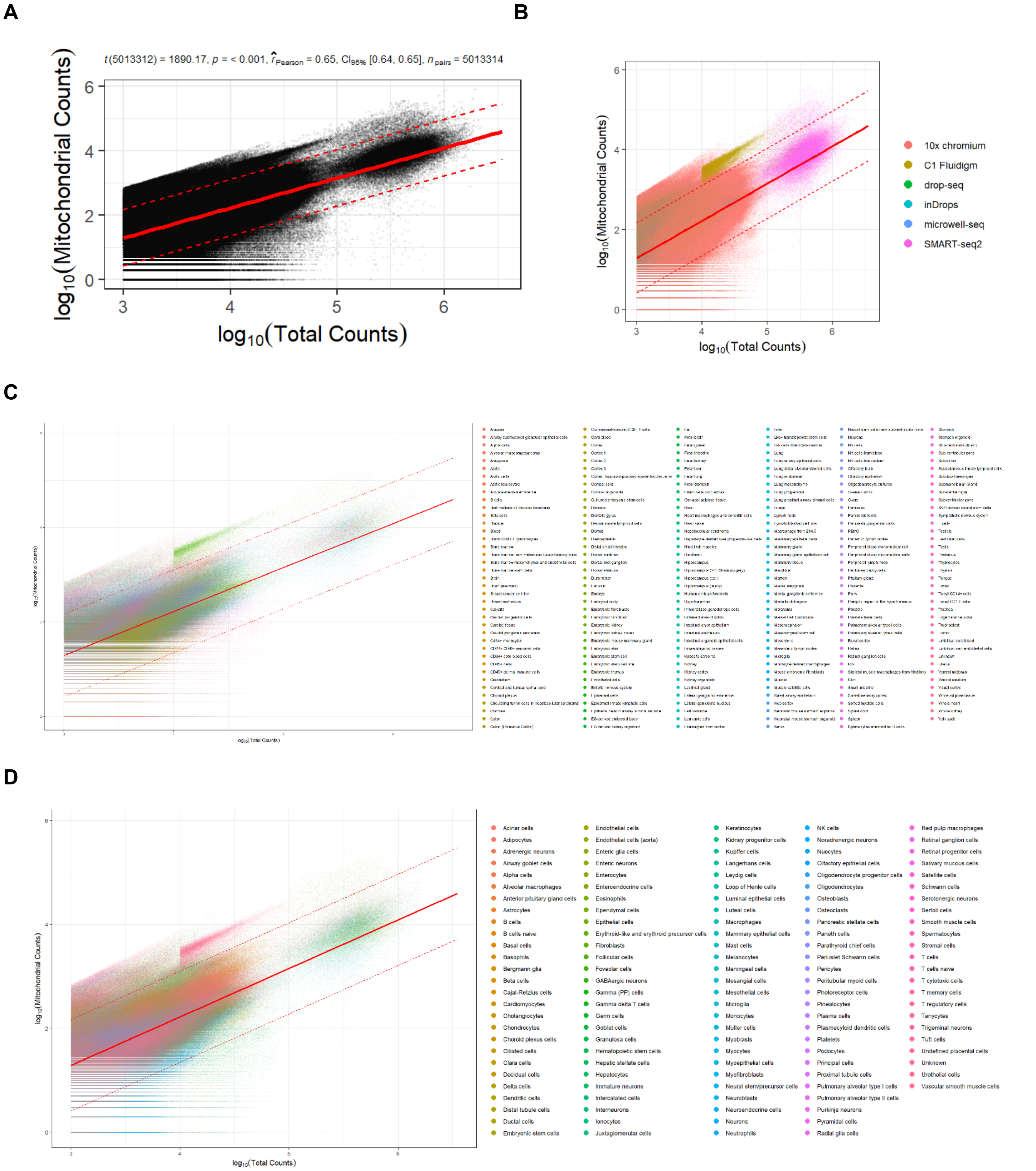


**Fig S2. Relationship between the total number of mitochondrial counts and the library size.** (A) Each dot represents a cell; the continuous red line represents the expectation and the dotted lines the lower and upper limit of the confidence interval of the prediction computed using OLS regression. (B) Same as in A colored by technology. (C) Same as in A colored by tissue. (D) Same as in A colored by cell-type.

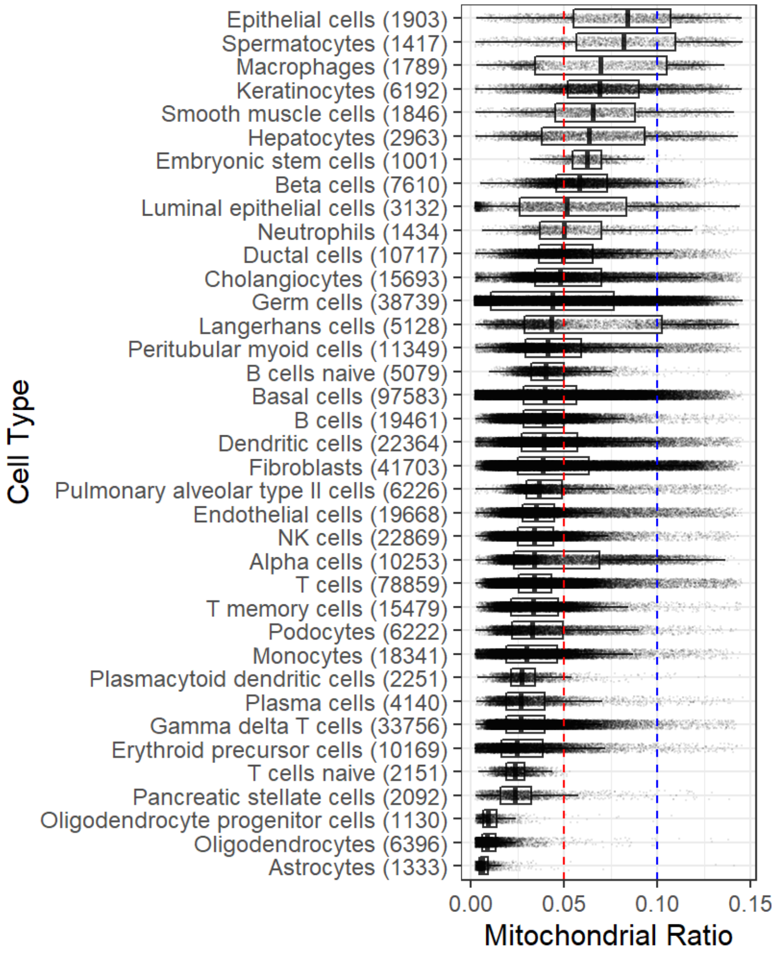


**Fig S3. Boxplots of the mitochondrial proportion across 37 human cell types.** In parenthesis, the number of cells in the stated cell type. Each dot represents a cell, the red line is the stablished 5% threshold, blue line is the 10% threshold for human cells proposed here.

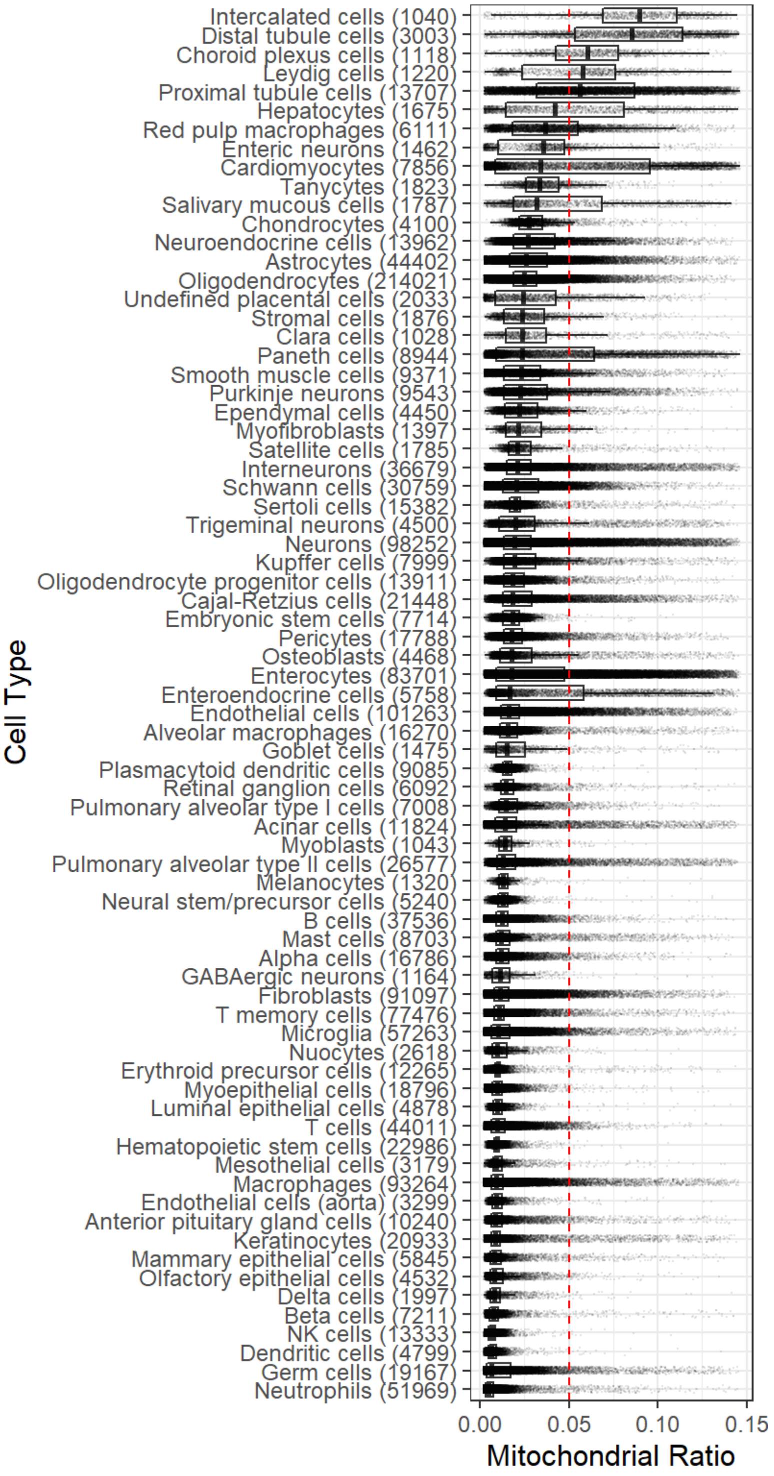


**Fig S4. Boxplots of the mitochondrial proportion across 75 mouse cell types.** In parenthesis, the number of cells in the stated cell type. Each dot represents a cell, the red line is the stablished 5% threshold.

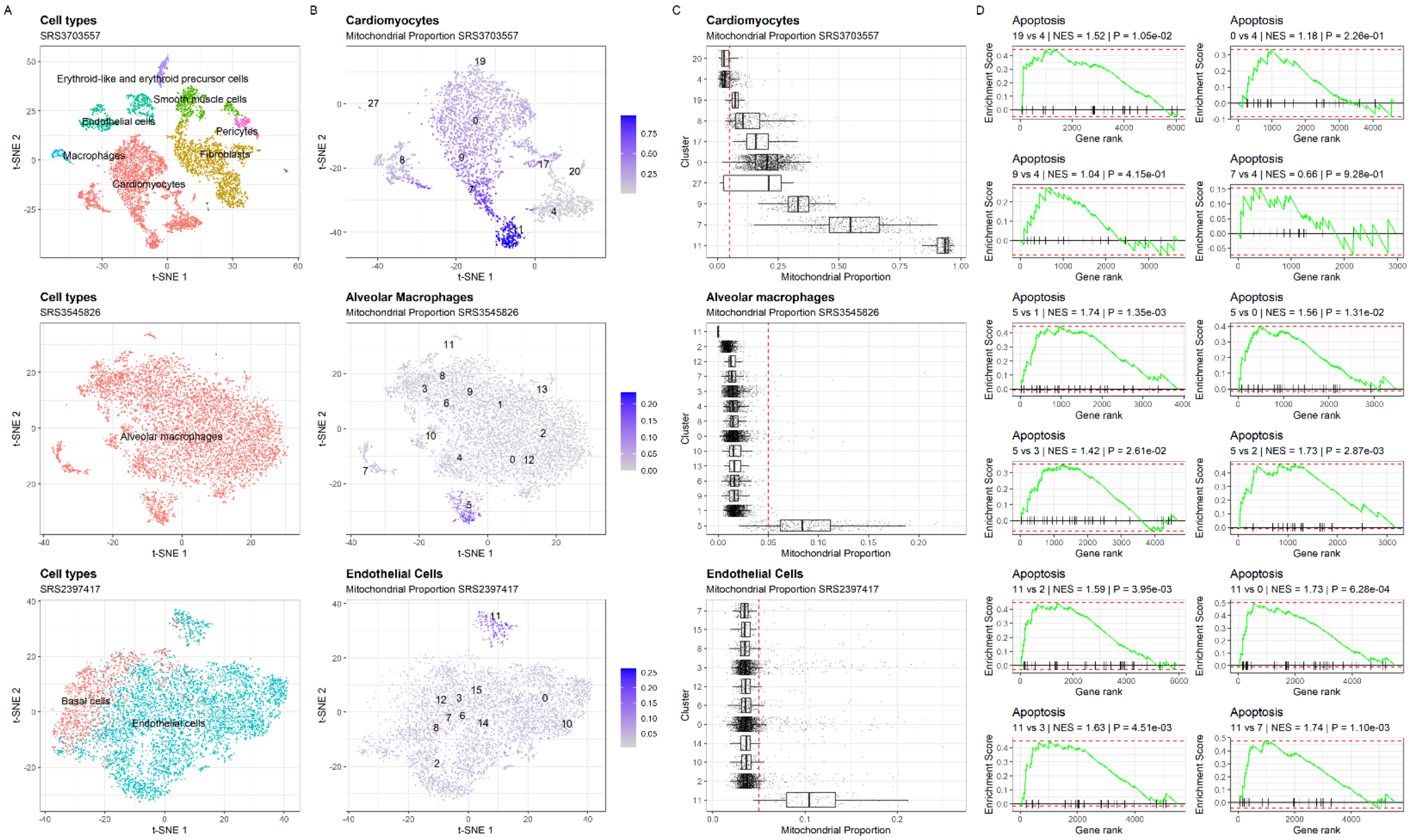


**Fig S5. Case examples showing the effect of omitting the mitochondrial proportion quality control filter in the analysis of single-cell RNA-seq data. (A)** t-SNE representation of all the cell populations included in the dataset generated by including the mitochondrial genes in the list of highly variable genes for principal component analysis (PCA). Each dot represents a cell and they are colored by cell-type. **(B)** t-SNE representation of cell-type used as example colored in function of the mitochondrial proportion in each cell. Clusters reported by the PanglaoDB are labeled. **(C)** Boxplot showing the distribution of the mitochondrial proportion across clusters. The red line is the early established 5% threshold. **(D)** Gene set enrichment analysis of the apoptosis pathway between clusters with high proportion of low-quality cells and others containing high quality cells.

**Table S1. Quantile values for mitochondrial proportion across 37 human cell types.** Min.: Minimum value, 1st Qu.: 25th percentile value, Median: 50th percentile value, Mean: Average value, 3rd Qu.: 75th percentile value, Max.: Maximum value.

| | Min. | 1st Qu. | Median | Mean | 3rd Qu. | Max. |
|---|---|---|---|---|---|---|
| **Alpha cells** | 0.003 | 0.023 | 0.034 | 0.046 | 0.069 | 0.144 |
| **Astrocytes** | 0.003 | 0.004 | 0.006 | 0.008 | 0.009 | 0.141 |
| **B cells** | 0.003 | 0.028 | 0.039 | 0.04 | 0.05 | 0.144 |
| **B cells naive** | 0.009 | 0.033 | 0.04 | 0.042 | 0.05 | 0.138 |
| **Basal cells** | 0.002 | 0.028 | 0.04 | 0.045 | 0.057 | 0.146 |
| **Beta cells** | 0.004 | 0.046 | 0.058 | 0.061 | 0.073 | 0.144 |
| **Cholangiocytes** | 0.002 | 0.034 | 0.048 | 0.055 | 0.07 | 0.145 |
| **Dendritic cells** | 0.002 | 0.027 | 0.039 | 0.045 | 0.057 | 0.146 |
| **Ductal cells** | 0.002 | 0.036 | 0.049 | 0.053 | 0.065 | 0.145 |
| **Embryonic stem cells** | 0.02 | 0.054 | 0.062 | 0.063 | 0.07 | 0.124 |
| **Endothelial cells** | 0.003 | 0.028 | 0.035 | 0.041 | 0.045 | 0.145 |
| **Epithelial cells** | 0.003 | 0.055 | 0.084 | 0.08 | 0.107 | 0.145 |
| **Erythroid precursor cells** | 0.002 | 0.016 | 0.025 | 0.03 | 0.039 | 0.145 |
| **Fibroblasts** | 0.002 | 0.025 | 0.039 | 0.047 | 0.063 | 0.145 |
| **Gamma delta T cells** | 0.003 | 0.019 | 0.026 | 0.031 | 0.039 | 0.142 |
| **Germ cells** | 0.002 | 0.011 | 0.044 | 0.048 | 0.077 | 0.146 |
| **Hepatocytes** | 0.003 | 0.038 | 0.063 | 0.067 | 0.093 | 0.143 |
| **Keratinocytes** | 0.002 | 0.052 | 0.069 | 0.071 | 0.09 | 0.145 |
| **Langerhans cells** | 0.003 | 0.029 | 0.043 | 0.061 | 0.102 | 0.144 |
| **Luminal epithelial cells** | 0.002 | 0.026 | 0.051 | 0.054 | 0.083 | 0.144 |
| **Macrophages** | 0.003 | 0.034 | 0.07 | 0.07 | 0.105 | 0.136 |
| **Monocytes** | 0.002 | 0.019 | 0.03 | 0.034 | 0.046 | 0.144 |
| **Neutrophils** | 0.006 | 0.037 | 0.05 | 0.057 | 0.07 | 0.143 |
| **NK cells** | 0.003 | 0.025 | 0.034 | 0.036 | 0.044 | 0.143 |
| **Oligodendrocyte progenitor cells** | 0.003 | 0.006 | 0.009 | 0.011 | 0.014 | 0.108 |
| **Oligodendrocytes** | 0.002 | 0.006 | 0.009 | 0.011 | 0.013 | 0.138 |
| **Pancreatic stellate cells** | 0.002 | 0.016 | 0.024 | 0.027 | 0.033 | 0.138 |
| **Peritubular myoid cells** | 0.002 | 0.029 | 0.041 | 0.048 | 0.059 | 0.145 |

| | | | | | | |
|---|---|---|---|---|---|---|
| **Plasma cells** | 0.003 | 0.019 | 0.027 | 0.032 | 0.039 | 0.146 |
| **Plasmacytoid dendritic cells** | 0.003 | 0.021 | 0.027 | 0.029 | 0.034 | 0.144 |
| **Podocytes** | 0.002 | 0.022 | 0.033 | 0.038 | 0.049 | 0.142 |
| **Pulmonary alveolar type II cells** | 0.002 | 0.03 | 0.037 | 0.044 | 0.049 | 0.144 |
| **Smooth muscle cells** | 0.003 | 0.045 | 0.066 | 0.068 | 0.088 | 0.141 |
| **Spermatocytes** | 0.002 | 0.056 | 0.082 | 0.081 | 0.109 | 0.146 |
| **T cells** | 0.002 | 0.026 | 0.034 | 0.036 | 0.043 | 0.145 |
| **T cells naive** | 0.004 | 0.019 | 0.024 | 0.024 | 0.029 | 0.053 |
| **T memory cells** | 0.003 | 0.021 | 0.033 | 0.036 | 0.047 | 0.145 |

**Table S2. Quantile values for mitochondrial proportion across 44 human tissues.** Min.: Minimum value, 1st Qu.: 25th percentile value, Median: 50th percentile value, Mean: Average value, 3rd Qu.: 75th percentile value, Max.: Maximum value.

| | Min. | 1st Qu. | Median | Mean | 3rd Qu. | Max. |
|---|---|---|---|---|---|---|
| **Adipose** | 0.011 | 0.089 | 0.104 | 0.101 | 0.115 | 0.145 |
| **B cells** | 0.003 | 0.024 | 0.032 | 0.034 | 0.04 | 0.137 |
| **Blood CD8+ T lymphocytes** | 0.009 | 0.033 | 0.039 | 0.041 | 0.047 | 0.139 |
| **Bone marrow** | 0.002 | 0.026 | 0.035 | 0.038 | 0.046 | 0.146 |
| **Brain** | 0.026 | 0.072 | 0.086 | 0.086 | 0.1 | 0.135 |
| **Breast epithelium** | 0.002 | 0.019 | 0.027 | 0.031 | 0.037 | 0.144 |
| **CD34+ cord blood cells** | 0.003 | 0.022 | 0.028 | 0.031 | 0.036 | 0.144 |
| **Colon** | 0.002 | 0.048 | 0.07 | 0.071 | 0.095 | 0.145 |
| **Colon (Ulcerative Colitis)** | 0.003 | 0.034 | 0.046 | 0.053 | 0.067 | 0.146 |
| **Decidua** | 0.002 | 0.012 | 0.019 | 0.028 | 0.035 | 0.141 |
| **Embryonic kidney** | 0.002 | 0.021 | 0.031 | 0.036 | 0.044 | 0.142 |
| **Embryonic kidney cortex** | 0.003 | 0.017 | 0.021 | 0.024 | 0.028 | 0.134 |
| **Embryonic stem cell line** | 0.002 | 0.02 | 0.03 | 0.037 | 0.045 | 0.146 |
| **ES-derived kidney organoid** | 0.003 | 0.034 | 0.049 | 0.052 | 0.065 | 0.14 |
| **Hepatocellular carcinoma** | 0.005 | 0.039 | 0.047 | 0.049 | 0.056 | 0.144 |
| **Hepatocyte-derived liver progenitor-like cells** | 0.002 | 0.029 | 0.035 | 0.037 | 0.042 | 0.133 |
| **Kaposi's sarcoma** | 0.02 | 0.054 | 0.062 | 0.063 | 0.07 | 0.124 |
| **Liver** | 0.003 | 0.065 | 0.088 | 0.087 | 0.111 | 0.144 |
| **Lung airway epithelial cells** | 0.002 | 0.007 | 0.014 | 0.018 | 0.026 | 0.144 |
| **Lung distal alveolar stromal cells** | 0.003 | 0.026 | 0.033 | 0.038 | 0.044 | 0.144 |
| **Lung proximal airway stromal cells** | 0.002 | 0.032 | 0.041 | 0.048 | 0.056 | 0.145 |
| **Lymph node** | 0.003 | 0.014 | 0.019 | 0.024 | 0.027 | 0.144 |
| **Lymphoblastoid cell line** | 0.003 | 0.028 | 0.04 | 0.04 | 0.049 | 0.146 |
| **Mammary gland** | 0.003 | 0.055 | 0.07 | 0.073 | 0.088 | 0.145 |
| **Melanoma** | 0.002 | 0.042 | 0.057 | 0.06 | 0.08 | 0.138 |
| **Monocyte-derived macrophages** | 0.007 | 0.085 | 0.104 | 0.1 | 0.119 | 0.144 |
| **Nasal airway epithelium** | 0.002 | 0.066 | 0.085 | 0.081 | 0.099 | 0.129 |
| **NK cells** | 0.003 | 0.015 | 0.019 | 0.021 | 0.024 | 0.139 |

| | | | | | | |
|---|---|---|---|---|---|---|
| **NK cells from blood** | 0.003 | 0.024 | 0.03 | 0.032 | 0.037 | 0.119 |
| **NK cells from spleen** | 0.003 | 0.027 | 0.034 | 0.035 | 0.042 | 0.111 |
| **Ovarian tumor** | 0.006 | 0.032 | 0.039 | 0.041 | 0.048 | 0.118 |
| **Pancreatic islets** | 0.002 | 0.024 | 0.037 | 0.042 | 0.056 | 0.144 |
| **Pancreatic progenitor cells** | 0.01 | 0.065 | 0.079 | 0.079 | 0.092 | 0.135 |
| **PBMC** | 0.003 | 0.022 | 0.029 | 0.034 | 0.04 | 0.146 |
| **Placenta** | 0.002 | 0.014 | 0.021 | 0.028 | 0.034 | 0.145 |
| **Prostate** | 0.003 | 0.033 | 0.043 | 0.048 | 0.058 | 0.145 |
| **Substantia nigra** | 0.002 | 0.006 | 0.009 | 0.012 | 0.014 | 0.141 |
| **T cells** | 0.002 | 0.028 | 0.037 | 0.038 | 0.046 | 0.145 |
| **Testicle** | 0.002 | 0.011 | 0.041 | 0.049 | 0.081 | 0.146 |
| **Testis** | 0.002 | 0.023 | 0.043 | 0.049 | 0.072 | 0.146 |
| **Tonsil** | 0.002 | 0.02 | 0.027 | 0.029 | 0.034 | 0.144 |
| **Tonsil CD14+ cells** | 0.002 | 0.023 | 0.031 | 0.033 | 0.041 | 0.138 |
| **Tonsil CD4 T cells** | 0.003 | 0.021 | 0.029 | 0.034 | 0.038 | 0.145 |
| **Umbilical vein endothelial cells** | 0.006 | 0.03 | 0.034 | 0.038 | 0.04 | 0.145 |

**Table S3. Quantile values for mitochondrial proportion across 74 mice cell types.** Min.: Minimum value, 1st Qu.: 25th percentile value, Median: 50th percentile value, Mean: Average value, 3rd Qu.: 75th percentile value, Max.: Maximum value.

| | **Min.** | **1st Qu.** | **Median** | **Mean** | **3rd Qu.** | **Max.** |
|---|---|---|---|---|---|---|
| **Acinar cells** | 0.002 | 0.009 | 0.014 | 0.022 | 0.021 | 0.145 |
| **Alpha cells** | 0.002 | 0.009 | 0.012 | 0.015 | 0.016 | 0.141 |
| **Alveolar macrophages** | 0.002 | 0.011 | 0.016 | 0.018 | 0.021 | 0.145 |
| **Anterior pituitary gland cells** | 0.002 | 0.006 | 0.009 | 0.013 | 0.013 | 0.144 |
| **Astrocytes** | 0.002 | 0.017 | 0.026 | 0.029 | 0.038 | 0.146 |
| **B cells** | 0.002 | 0.009 | 0.012 | 0.013 | 0.016 | 0.143 |
| **Beta cells** | 0.002 | 0.005 | 0.007 | 0.01 | 0.01 | 0.143 |
| **Cajal-Retzius cells** | 0.002 | 0.013 | 0.018 | 0.025 | 0.029 | 0.145 |
| **Cardiomyocytes** | 0.002 | 0.009 | 0.034 | 0.052 | 0.096 | 0.146 |
| **Chondrocytes** | 0.004 | 0.022 | 0.028 | 0.031 | 0.035 | 0.133 |
| **Choroid plexus cells** | 0.003 | 0.043 | 0.06 | 0.061 | 0.078 | 0.138 |
| **Clara cells** | 0.002 | 0.014 | 0.024 | 0.033 | 0.037 | 0.145 |
| **Delta cells** | 0.002 | 0.006 | 0.008 | 0.011 | 0.012 | 0.142 |
| **Dendritic cells** | 0.002 | 0.005 | 0.007 | 0.008 | 0.009 | 0.135 |
| **Distal tubule cells** | 0.003 | 0.053 | 0.085 | 0.083 | 0.114 | 0.146 |
| **Embryonic stem cells** | 0.002 | 0.013 | 0.018 | 0.018 | 0.022 | 0.108 |
| **Endothelial cells** | 0.002 | 0.012 | 0.016 | 0.02 | 0.022 | 0.144 |
| **Endothelial cells (aorta)** | 0.003 | 0.007 | 0.009 | 0.011 | 0.012 | 0.13 |
| **Enteric neurons** | 0.002 | 0.01 | 0.036 | 0.036 | 0.048 | 0.141 |
| **Enterocytes** | 0.002 | 0.009 | 0.018 | 0.032 | 0.048 | 0.146 |
| **Enteroendocrine cells** | 0.002 | 0.009 | 0.017 | 0.037 | 0.058 | 0.146 |
| **Ependymal cells** | 0.003 | 0.014 | 0.022 | 0.026 | 0.032 | 0.142 |
| **Erythroid precursor cells** | 0.002 | 0.008 | 0.01 | 0.011 | 0.011 | 0.144 |
| **Fibroblasts** | 0.002 | 0.008 | 0.011 | 0.015 | 0.016 | 0.143 |
| **GABAergic neurons** | 0.002 | 0.007 | 0.012 | 0.014 | 0.017 | 0.131 |
| **Germ cells** | 0.002 | 0.004 | 0.006 | 0.014 | 0.017 | 0.145 |
| **Goblet cells** | 0.002 | 0.009 | 0.015 | 0.022 | 0.025 | 0.144 |
| **Hematopoietic stem cells** | 0.003 | 0.008 | 0.009 | 0.01 | 0.011 | 0.106 |

| | | | | | | |
|---|---|---|---|---|---|---|
| **Hepatocytes** | 0.002 | 0.015 | 0.042 | 0.05 | 0.081 | 0.145 |
| **Intercalated cells** | 0.002 | 0.069 | 0.089 | 0.086 | 0.111 | 0.145 |
| **Interneurons** | 0.002 | 0.015 | 0.021 | 0.026 | 0.028 | 0.146 |
| **Keratinocytes** | 0.002 | 0.006 | 0.008 | 0.012 | 0.011 | 0.144 |
| **Kupffer cells** | 0.002 | 0.013 | 0.019 | 0.026 | 0.031 | 0.145 |
| **Leydig cells** | 0.003 | 0.024 | 0.058 | 0.054 | 0.076 | 0.141 |
| **Luminal epithelial cells** | 0.002 | 0.008 | 0.01 | 0.011 | 0.012 | 0.141 |
| **Macrophages** | 0.002 | 0.006 | 0.009 | 0.012 | 0.013 | 0.145 |
| **Mammary epithelial cells** | 0.002 | 0.006 | 0.008 | 0.012 | 0.012 | 0.137 |
| **Mast cells** | 0.002 | 0.009 | 0.012 | 0.018 | 0.017 | 0.142 |
| **Melanocytes** | 0.003 | 0.011 | 0.013 | 0.014 | 0.016 | 0.124 |
| **Mesothelial cells** | 0.002 | 0.007 | 0.009 | 0.012 | 0.013 | 0.138 |
| **Microglia** | 0.002 | 0.006 | 0.01 | 0.014 | 0.017 | 0.144 |
| **Myoblasts** | 0.003 | 0.011 | 0.014 | 0.016 | 0.018 | 0.094 |
| **Myoepithelial cells** | 0.002 | 0.007 | 0.01 | 0.011 | 0.013 | 0.128 |
| **Myofibroblasts** | 0.003 | 0.014 | 0.022 | 0.027 | 0.034 | 0.145 |
| **Neural stem/precursor cells** | 0.002 | 0.01 | 0.013 | 0.013 | 0.016 | 0.131 |
| **Neuroendocrine cells** | 0.002 | 0.019 | 0.027 | 0.033 | 0.042 | 0.142 |
| **Neurons** | 0.002 | 0.013 | 0.019 | 0.025 | 0.029 | 0.146 |
| **Neutrophils** | 0.002 | 0.003 | 0.005 | 0.006 | 0.008 | 0.145 |
| **NK cells** | 0.002 | 0.005 | 0.007 | 0.007 | 0.009 | 0.107 |
| **Nuocytes** | 0.002 | 0.007 | 0.01 | 0.013 | 0.015 | 0.112 |
| **Olfactory epithelial cells** | 0.002 | 0.006 | 0.008 | 0.012 | 0.013 | 0.134 |
| **Oligodendrocyte progenitor cells** | 0.002 | 0.013 | 0.019 | 0.021 | 0.025 | 0.142 |
| **Oligodendrocytes** | 0.002 | 0.019 | 0.025 | 0.026 | 0.032 | 0.143 |
| **Osteoblasts** | 0.003 | 0.012 | 0.018 | 0.026 | 0.029 | 0.145 |
| **Paneth cells** | 0.002 | 0.009 | 0.024 | 0.04 | 0.064 | 0.146 |
| **Pericytes** | 0.003 | 0.014 | 0.018 | 0.021 | 0.024 | 0.14 |
| **Plasmacytoid dendritic cells** | 0.003 | 0.013 | 0.015 | 0.016 | 0.018 | 0.122 |
| **Proximal tubule cells** | 0.002 | 0.032 | 0.056 | 0.061 | 0.087 | 0.146 |

| | | | | | | |
|---|---|---|---|---|---|---|
| **Pulmonary alveolar type I cells** | 0.002 | 0.011 | 0.014 | 0.019 | 0.021 | 0.134 |
| **Pulmonary alveolar type II cells** | 0.002 | 0.01 | 0.013 | 0.019 | 0.02 | 0.145 |
| **Purkinje neurons** | 0.002 | 0.014 | 0.022 | 0.03 | 0.038 | 0.145 |
| **Red pulp macrophages** | 0.002 | 0.019 | 0.037 | 0.04 | 0.055 | 0.143 |
| **Retinal ganglion cells** | 0.003 | 0.012 | 0.015 | 0.018 | 0.019 | 0.143 |
| **Salivary mucous cells** | 0.002 | 0.019 | 0.032 | 0.046 | 0.068 | 0.144 |
| **Satellite cells** | 0.005 | 0.016 | 0.021 | 0.028 | 0.029 | 0.137 |
| **Schwann cells** | 0.002 | 0.013 | 0.021 | 0.024 | 0.033 | 0.146 |
| **Sertoli cells** | 0.002 | 0.017 | 0.02 | 0.021 | 0.023 | 0.143 |
| **Smooth muscle cells** | 0.002 | 0.013 | 0.023 | 0.027 | 0.034 | 0.143 |
| **Stromal cells** | 0.003 | 0.014 | 0.024 | 0.028 | 0.036 | 0.141 |
| **T cells** | 0.002 | 0.006 | 0.01 | 0.012 | 0.014 | 0.136 |
| **T memory cells** | 0.002 | 0.008 | 0.011 | 0.012 | 0.014 | 0.142 |
| **Tanycytes** | 0.003 | 0.026 | 0.034 | 0.037 | 0.044 | 0.136 |
| **Trigeminal neurons** | 0.002 | 0.011 | 0.02 | 0.03 | 0.031 | 0.145 |
| **Undefined placental cells** | 0.002 | 0.009 | 0.024 | 0.031 | 0.043 | 0.138 |

**Table S4. Quantile values for mitochondrial proportion across 121 mouse tissues.** Min.: Minimum value, 1st Qu.: 25th percentile value, Median: 50th percentile value, Mean: Average value, 3rd Qu.: 75th percentile value, Max.: Maximum value.

| | Min. | 1st Qu. | Median | Mean | 3rd Qu. | Max. |
|---|---|---|---|---|---|---|
| **Amygdala** | 0.003 | 0.016 | 0.021 | 0.022 | 0.026 | 0.139 |
| **Aorta** | 0.002 | 0.007 | 0.01 | 0.012 | 0.013 | 0.145 |
| **Aortic leukocytes** | 0.002 | 0.007 | 0.01 | 0.013 | 0.015 | 0.141 |
| **Bed nucleus of the stria terminalis** | 0.002 | 0.004 | 0.006 | 0.009 | 0.011 | 0.123 |
| **Bladder** | 0.003 | 0.009 | 0.011 | 0.014 | 0.013 | 0.134 |
| **Bone marrow** | 0.002 | 0.007 | 0.011 | 0.013 | 0.016 | 0.145 |
| **Bone marrow from melanoma tumor-bearing mice** | 0.002 | 0.003 | 0.004 | 0.005 | 0.006 | 0.137 |
| **Bone marrow mesenchymal and endothelial cells** | 0.003 | 0.011 | 0.017 | 0.026 | 0.029 | 0.145 |
| **Brain** | 0.002 | 0.005 | 0.008 | 0.009 | 0.011 | 0.136 |
| **Cardiac progenitor cells** | 0.002 | 0.009 | 0.012 | 0.017 | 0.017 | 0.135 |
| **Cardiac tissue** | 0.002 | 0.005 | 0.009 | 0.02 | 0.022 | 0.141 |
| **CD45+ cells** | 0.002 | 0.003 | 0.004 | 0.005 | 0.006 | 0.075 |
| **CD45+ dermal immune cells** | 0.003 | 0.008 | 0.011 | 0.014 | 0.016 | 0.094 |
| **Cerebellum** | 0.003 | 0.021 | 0.028 | 0.031 | 0.036 | 0.136 |
| **Choroid plexus** | 0.002 | 0.006 | 0.008 | 0.009 | 0.01 | 0.135 |
| **Colon** | 0.002 | 0.01 | 0.015 | 0.021 | 0.024 | 0.145 |
| **Commensal-specific CD8+ T cells** | 0.002 | 0.006 | 0.008 | 0.01 | 0.011 | 0.115 |
| **Cortex** | 0.003 | 0.015 | 0.022 | 0.028 | 0.032 | 0.146 |
| **Cortex, hippocampus and subventricular zone** | 0.002 | 0.013 | 0.017 | 0.021 | 0.023 | 0.145 |
| **Dentate gyrus** | 0.003 | 0.008 | 0.012 | 0.016 | 0.02 | 0.133 |
| **Dermal innate lymphoid cells** | 0.003 | 0.009 | 0.012 | 0.013 | 0.015 | 0.107 |
| **Dermis** | 0.003 | 0.012 | 0.017 | 0.021 | 0.025 | 0.141 |
| **Distal small intestine** | 0.002 | 0.036 | 0.053 | 0.059 | 0.077 | 0.146 |
| **Dorsal midbrain** | 0.002 | 0.015 | 0.021 | 0.024 | 0.031 | 0.144 |
| **Dorsal root ganglion** | 0.002 | 0.005 | 0.007 | 0.01 | 0.013 | 0.094 |
| **Dorsal striatum** | 0.002 | 0.009 | 0.015 | 0.018 | 0.021 | 0.136 |
| **Dura mater** | 0.002 | 0.005 | 0.008 | 0.009 | 0.012 | 0.096 |

| | | | | | | |
|---|---|---|---|---|---|---|
| **Ear skin** | 0.005 | 0.028 | 0.034 | 0.037 | 0.042 | 0.128 |
| **Embryo** | 0.002 | 0.018 | 0.029 | 0.035 | 0.048 | 0.139 |
| **Embryonic fibroblasts** | 0.002 | 0.013 | 0.018 | 0.019 | 0.022 | 0.127 |
| **Embryonic hindbrain** | 0.003 | 0.012 | 0.017 | 0.018 | 0.021 | 0.124 |
| **Embryonic kidney** | 0.003 | 0.014 | 0.017 | 0.02 | 0.021 | 0.139 |
| **Embryonic skin** | 0.002 | 0.01 | 0.012 | 0.017 | 0.015 | 0.135 |
| **Enteric nervous system** | 0.003 | 0.015 | 0.021 | 0.024 | 0.03 | 0.146 |
| **Epidermal cells** | 0.002 | 0.009 | 0.012 | 0.015 | 0.015 | 0.14 |
| **Epidermal innate lymphoid cells** | 0.002 | 0.01 | 0.012 | 0.014 | 0.015 | 0.101 |
| **Gonadal adipose tissue** | 0.002 | 0.005 | 0.007 | 0.007 | 0.009 | 0.042 |
| **Heart** | 0.002 | 0.016 | 0.026 | 0.033 | 0.04 | 0.143 |
| **Heart macrophages and dendritic cells** | 0.002 | 0.005 | 0.007 | 0.01 | 0.011 | 0.135 |
| **Hind limb muscles** | 0.002 | 0.009 | 0.013 | 0.015 | 0.017 | 0.097 |
| **Hindbrain** | 0.003 | 0.014 | 0.019 | 0.022 | 0.026 | 0.146 |
| **Hippocampus** | 0.002 | 0.014 | 0.021 | 0.025 | 0.03 | 0.144 |
| **Hippocampus (ca1)** | 0.002 | 0.009 | 0.016 | 0.021 | 0.025 | 0.144 |
| **Hypothalamus** | 0.002 | 0.016 | 0.021 | 0.027 | 0.03 | 0.146 |
| **Intestinal crypt epithelium** | 0.003 | 0.009 | 0.011 | 0.017 | 0.015 | 0.146 |
| **Intestinal epithelium** | 0.003 | 0.017 | 0.035 | 0.043 | 0.062 | 0.144 |
| **Intestinal organoid epithelial cells** | 0.002 | 0.008 | 0.013 | 0.022 | 0.028 | 0.145 |
| **Intraembryonic tissues** | 0.002 | 0.01 | 0.013 | 0.02 | 0.017 | 0.142 |
| **Kaposi's sarcoma** | 0.003 | 0.013 | 0.015 | 0.016 | 0.018 | 0.116 |
| **Kidney** | 0.003 | 0.015 | 0.019 | 0.035 | 0.037 | 0.145 |
| **Left Ventricle** | 0.003 | 0.012 | 0.017 | 0.024 | 0.027 | 0.144 |
| **Leukemic cells** | 0.002 | 0.007 | 0.009 | 0.009 | 0.011 | 0.065 |
| **Leukocytes from aortas** | 0.002 | 0.007 | 0.009 | 0.011 | 0.012 | 0.1 |
| **Liver** | 0.002 | 0.011 | 0.017 | 0.025 | 0.029 | 0.145 |
| **Lks+ hematopoietic stem cells** | 0.003 | 0.009 | 0.01 | 0.01 | 0.012 | 0.046 |
| **Lsk cells from bone marrow** | 0.003 | 0.008 | 0.01 | 0.01 | 0.011 | 0.134 |
| **Lung** | 0.002 | 0.01 | 0.014 | 0.017 | 0.019 | 0.145 |
| **Lung endoderm** | 0.002 | 0.01 | 0.012 | 0.014 | 0.015 | 0.117 |
| **Lung mesenchyme** | 0.002 | 0.003 | 0.004 | 0.005 | 0.005 | 0.052 |

| | | | | | | |
|---|---|---|---|---|---|---|
| **Macrophage from E14.5** | 0.002 | 0.007 | 0.008 | 0.01 | 0.011 | 0.144 |
| **Mammary epithelial cells** | 0.002 | 0.007 | 0.01 | 0.011 | 0.013 | 0.137 |
| **Mammary gland epithelium cell** | 0.002 | 0.005 | 0.007 | 0.008 | 0.009 | 0.055 |
| **Mammary tissue** | 0.002 | 0.01 | 0.012 | 0.014 | 0.016 | 0.141 |
| **Marrow** | 0.002 | 0.005 | 0.009 | 0.01 | 0.013 | 0.143 |
| **Medulla oblongata** | 0.002 | 0.021 | 0.026 | 0.028 | 0.033 | 0.142 |
| **Mesenteric** | 0.002 | 0.01 | 0.013 | 0.015 | 0.017 | 0.116 |
| **Mesenteric lymph nodes** | 0.002 | 0.009 | 0.013 | 0.02 | 0.027 | 0.143 |
| **Microglia** | 0.002 | 0.011 | 0.017 | 0.024 | 0.025 | 0.144 |
| **Mouse embryonic fibroblasts** | 0.003 | 0.014 | 0.018 | 0.02 | 0.023 | 0.082 |
| **Muscle** | 0.002 | 0.008 | 0.012 | 0.015 | 0.016 | 0.137 |
| **Muscle satellite cells** | 0.003 | 0.015 | 0.019 | 0.025 | 0.025 | 0.146 |
| **Neo-cortex** | 0.002 | 0.013 | 0.018 | 0.022 | 0.025 | 0.144 |
| **Neonatal mouse stomach explants** | 0.002 | 0.004 | 0.005 | 0.008 | 0.006 | 0.143 |
| **Nerve** | 0.002 | 0.021 | 0.032 | 0.047 | 0.07 | 0.145 |
| **NK cells from blood** | 0.003 | 0.005 | 0.007 | 0.007 | 0.009 | 0.024 |
| **NK cells from spleen** | 0.002 | 0.004 | 0.005 | 0.006 | 0.007 | 0.027 |
| **Olfactory bulb** | 0.003 | 0.019 | 0.029 | 0.033 | 0.042 | 0.145 |
| **Olfactory epithelium** | 0.002 | 0.006 | 0.008 | 0.012 | 0.014 | 0.125 |
| **Pancreas** | 0.002 | 0.007 | 0.01 | 0.011 | 0.014 | 0.141 |
| **Pancreatic islets** | 0.002 | 0.01 | 0.015 | 0.024 | 0.027 | 0.146 |
| **Pancreatic progenitor cells** | 0.002 | 0.008 | 0.011 | 0.013 | 0.015 | 0.135 |
| **Periaortic lymph nodes** | 0.003 | 0.008 | 0.011 | 0.017 | 0.021 | 0.131 |
| **Peripheral lymph node** | 0.005 | 0.016 | 0.019 | 0.02 | 0.023 | 0.052 |
| **Peritoneal cavity cells** | 0.002 | 0.01 | 0.012 | 0.013 | 0.015 | 0.117 |
| **Pituitary gland** | 0.002 | 0.007 | 0.009 | 0.013 | 0.013 | 0.144 |
| **Pons** | 0.002 | 0.022 | 0.027 | 0.028 | 0.033 | 0.144 |
| **Preoptic region of the hypothalamus** | 0.002 | 0.017 | 0.024 | 0.034 | 0.041 | 0.146 |
| **Prostate basal cells** | 0.002 | 0.006 | 0.008 | 0.01 | 0.01 | 0.144 |
| **Pulmonary alveolar type i cells** | 0.002 | 0.011 | 0.014 | 0.018 | 0.02 | 0.134 |
| **Pulmonary alveolar type ii cells** | 0.003 | 0.019 | 0.025 | 0.032 | 0.036 | 0.144 |
| **Retinal ganglion cells** | 0.003 | 0.012 | 0.014 | 0.017 | 0.019 | 0.143 |

| | | | | | | |
|---|---|---|---|---|---|---|
| **Skeletal muscle macrophages from hindlimb** | 0.002 | 0.007 | 0.009 | 0.01 | 0.012 | 0.108 |
| **Skin** | 0.002 | 0.007 | 0.009 | 0.011 | 0.012 | 0.139 |
| **Small intestine** | 0.002 | 0.006 | 0.008 | 0.012 | 0.011 | 0.146 |
| **Somatosensory cortex** | 0.002 | 0.009 | 0.013 | 0.022 | 0.023 | 0.145 |
| **Sorted myeloid cells** | 0.002 | 0.005 | 0.007 | 0.008 | 0.009 | 0.085 |
| **Spinal cord** | 0.002 | 0.014 | 0.02 | 0.021 | 0.026 | 0.134 |
| **Spleen** | 0.002 | 0.008 | 0.011 | 0.014 | 0.015 | 0.143 |
| **Splenocytes enriched for B cells** | 0.003 | 0.007 | 0.009 | 0.009 | 0.011 | 0.036 |
| **Stria terminalis (brain)** | 0.002 | 0.003 | 0.005 | 0.008 | 0.01 | 0.095 |
| **Sub-ventricular zone** | 0.002 | 0.007 | 0.011 | 0.012 | 0.015 | 0.089 |
| **Subcortex** | 0.002 | 0.016 | 0.025 | 0.031 | 0.041 | 0.137 |
| **Subcutaneous innate lymphoid cells** | 0.002 | 0.005 | 0.008 | 0.01 | 0.011 | 0.118 |
| **Subdural meninges** | 0.002 | 0.005 | 0.008 | 0.008 | 0.011 | 0.137 |
| **Submandibular Gland** | 0.002 | 0.017 | 0.027 | 0.04 | 0.054 | 0.144 |
| **Subventricular zone** | 0.002 | 0.012 | 0.016 | 0.019 | 0.021 | 0.123 |
| **Sympathetic nervous system** | 0.002 | 0.004 | 0.005 | 0.012 | 0.009 | 0.144 |
| **T cells** | 0.002 | 0.004 | 0.006 | 0.007 | 0.008 | 0.127 |
| **Testis** | 0.002 | 0.007 | 0.015 | 0.018 | 0.022 | 0.145 |
| **Thalamus** | 0.003 | 0.019 | 0.025 | 0.028 | 0.034 | 0.146 |
| **Thymocytes** | 0.003 | 0.011 | 0.013 | 0.014 | 0.016 | 0.142 |
| **Thymus** | 0.002 | 0.009 | 0.014 | 0.016 | 0.022 | 0.118 |
| **Tongue** | 0.002 | 0.006 | 0.008 | 0.012 | 0.01 | 0.144 |
| **Trachea** | 0.006 | 0.022 | 0.028 | 0.032 | 0.036 | 0.145 |
| **Ventral midbrain** | 0.003 | 0.016 | 0.024 | 0.025 | 0.032 | 0.144 |
| **Ventral striatum** | 0.003 | 0.018 | 0.024 | 0.026 | 0.031 | 0.143 |
| **Visual cortex** | 0.003 | 0.012 | 0.015 | 0.017 | 0.019 | 0.122 |
| **White adipose tissue** | 0.005 | 0.009 | 0.011 | 0.012 | 0.014 | 0.07 |
| **Whole heart** | 0.002 | 0.013 | 0.022 | 0.051 | 0.097 | 0.146 |
| **Whole kidney** | 0.002 | 0.032 | 0.059 | 0.063 | 0.092 | 0.146 |
| **Yolk sack** | 0.002 | 0.011 | 0.014 | 0.02 | 0.02 | 0.133 |

**Table S5. Differences in mitochondrial proportion between human and mice cell types.** Mean Hsa: Mean mitochondrial proportion for human cells. Mean Mmu: Mean mitochondrial proportion for mice cells. Lower CI: Lower limit of the 95% CI for the difference in the means between human and mice cells. Upper CI: Upper limit of the 95% CI for the difference in the means between human and mice cells. P-value: the p-value for the T test. P-adj: the p-value for the T test after multiple testing correction (FDR). Hsa > Mmu: A logical vector displaying if the mean value for humans is greater than the mean value for mice.

| | Mean Hsa | Mean Mmu | Lower CI | Upper CI | P-value | P-adj | Hsa > Mmu |
|---|---|---|---|---|---|---|---|
| **NK cells** | 0.039 | 0.007 | 0.032 | 0.033 | 0.000 | 0.000 | TRUE |
| **T cells** | 0.040 | 0.011 | 0.029 | 0.029 | 0.000 | 0.000 | TRUE |
| **Plasmacytoid dendritic cells** | 0.032 | 0.016 | 0.015 | 0.018 | 0.000 | 0.000 | TRUE |
| **Dendritic cells** | 0.050 | 0.007 | 0.042 | 0.043 | 0.000 | 0.000 | TRUE |
| **Erythroid-like and erythroid precursor cells** | 0.015 | 0.005 | 0.010 | 0.011 | 0.000 | 0.000 | TRUE |
| **B cells** | 0.043 | 0.013 | 0.029 | 0.030 | 0.000 | 0.000 | TRUE |
| **Endothelial cells** | 0.053 | 0.020 | 0.032 | 0.034 | 0.000 | 0.000 | TRUE |
| **Luminal epithelial cells** | 0.107 | 0.011 | 0.090 | 0.102 | 0.000 | 0.000 | TRUE |
| **Keratinocytes** | 0.090 | 0.014 | 0.074 | 0.077 | 0.000 | 0.000 | TRUE |
| **Hepatocytes** | 0.658 | 0.210 | 0.438 | 0.458 | 0.000 | 0.000 | TRUE |
| **Pulmonary alveolar type II cells** | 0.051 | 0.022 | 0.028 | 0.030 | 0.000 | 0.000 | TRUE |
| **Smooth muscle cells** | 0.080 | 0.027 | 0.050 | 0.055 | 6.374e-315 | 8.606e-315 | TRUE |
| **Fibroblasts** | 0.055 | 0.015 | 0.039 | 0.040 | 0.000 | 0.000 | TRUE |
| **Germ cells** | 0.060 | 0.007 | 0.052 | 0.054 | 0.000 | 0.000 | TRUE |
| **Macrophages** | 0.095 | 0.012 | 0.080 | 0.087 | 0.000 | 0.000 | TRUE |
| **T memory cells** | 0.036 | 0.012 | 0.024 | 0.025 | 0.000 | 0.000 | TRUE |
| **Goblet cells** | 0.460 | 0.026 | 0.421 | 0.446 | 0.000 | 0.000 | TRUE |
| **Acinar cells** | 0.047 | 0.053 | -0.011 | -0.001 | 0.028 | 0.028 | FALSE |
| **Beta cells** | 0.072 | 0.039 | 0.031 | 0.036 | 0.000 | 0.000 | TRUE |
| **Alpha cells** | 0.064 | 0.014 | 0.048 | 0.051 | 0.000 | 0.000 | TRUE |
| **Enteroendocrine cells** | 0.249 | 0.098 | 0.137 | 0.166 | 0.000 | 0.000 | TRUE |
| **Neutrophils** | 0.062 | 0.005 | 0.054 | 0.058 | 0.000 | 0.000 | TRUE |
| **Enterocytes** | 0.245 | 0.049 | 0.187 | 0.205 | 0.000 | 0.000 | TRUE |
| **Embryonic stem cells** | 0.064 | 0.018 | 0.045 | 0.047 | 0.000 | 0.000 | TRUE |
| **Oligodendrocytes** | 0.011 | 0.025 | -0.015 | -0.014 | 0.000 | 0.000 | FALSE |
| **Astrocytes** | 0.007 | 0.025 | -0.018 | -0.017 | 0.000 | 0.000 | FALSE |
| **Oligodendrocyte progenitor cells** | 0.011 | 0.018 | -0.008 | -0.007 | 0.000 | 0.000 | FALSE |

**Table S6. Differences in mitochondrial proportion between human and mice tissues.** Mean Hsa: Mean mitochondrial proportion for human cells. Mean Mmu: Mean mitochondrial proportion for mice cells. Lower CI: Lower limit of the 95% CI for the difference in the means between human and mice cells. Upper CI: Upper limit of the 95% CI for the difference in the means between human and mice cells. P-value: the p-value for the T test. P-adj: the p-value for the T test after multiple testing correction (FDR). Hsa > Mmu: A logical vector displaying if the mean value for humans is greater than the mean value for mice.

| | Mean Hsa | Mean Mmu | Lower CI | Upper CI | P-value | P-adj | Hsa > Mmu |
|---|---|---|---|---|---|---|---|
| **Bone marrow** | 0.034 | 0.012 | 0.022 | 0.022 | 0.000 | 0.000 | TRUE |
| **Testis** | 0.108 | 0.015 | 0.091 | 0.093 | 0.000 | 0.000 | TRUE |
| **T cells** | 0.036 | 0.007 | 0.029 | 0.030 | 0.000 | 0.000 | TRUE |
| **Brain** | 0.095 | 0.009 | 0.084 | 0.089 | 0.000 | 0.000 | TRUE |
| **Pancreatic islets** | 0.077 | 0.039 | 0.036 | 0.039 | 0.000 | 0.000 | TRUE |
| **Colon** | 0.284 | 0.023 | 0.258 | 0.263 | 0.000 | 0.000 | TRUE |
| **Embryonic kidney** | 0.052 | 0.022 | 0.029 | 0.031 | 0.000 | 0.000 | TRUE |
| **Liver** | 0.575 | 0.048 | 0.521 | 0.531 | 0.000 | 0.000 | TRUE |
| **NK cells from spleen** | 0.035 | 0.005 | 0.029 | 0.030 | 0.000 | 0.000 | TRUE |
| **NK cells from blood** | 0.031 | 0.007 | 0.024 | 0.024 | 0.000 | 0.000 | TRUE |
| **Pancreatic progenitor cells** | 0.081 | 0.014 | 0.065 | 0.068 | 0.000 | 0.000 | TRUE |
| **Spleen** | 0.041 | 0.014 | 0.026 | 0.028 | 0.000 | 0.000 | TRUE |
| **Mammary gland** | 0.084 | 0.004 | 0.079 | 0.080 | 0.000 | 0.000 | TRUE |
| **Kaposi's sarcoma** | 0.043 | 0.016 | 0.026 | 0.027 | 0.000 | 0.000 | TRUE |
